# Plastid-specific RsmD methyltransferase and ribosome maturation factor RimM are crucial for 16S rRNA maturation and proteostasis

**DOI:** 10.1101/2022.03.07.483362

**Authors:** Kaiwei Liu, Keun Pyo Lee, Jianli Duan, Eun Yu Kim, Rahul Mohan Singh, Minghui Di, Zhuoling Meng, Chanhong Kim

## Abstract

Chloroplast pre-ribosomal RNA (rRNA) undergoes maturation, which is critical for ribosome assembly. While the central and auxiliary factors in rRNA maturation have been elucidated in bacteria, their mode of action remains largely unexplored in chloroplasts. We now reveal chloroplast-specific factors involved in 16S rRNA maturation, RsmD methyltransferase (AtRsmD) and ribosome maturation factor RimM-like protein (AtRimM) in *Arabidopsis thaliana.* A forward genetic screen aimed to find suppressors of the Arabidopsis *yellow variegated 2* (*var2*) mutant defective in photosystem II (PSII) quality control found a causal nonsense mutation in *AtRsmD*. The substantially impaired 16S rRNA maturation and translation due to the mutation rescued the leaf variegation phenotype by lowering the levels of PSII core proteins in *var2*. The subsequent co-immunoprecipitation coupled with mass spectrometry analyses and bimolecular fluorescence complementation assay found that AtRsmD interacts with AtRimM. Consistent with their interaction, loss of AtRimM also considerably impairs 16S rRNA maturation, with less methylation in m^2^G915 in 16S rRNA catalyzed by AtRsmD. The *atrimM* mutation also rescued *var2* mutant phenotypes, corroborating the functional interplay between AtRsmD and AtRimM towards 16S rRNA maturation and chloroplast proteostasis.

## INTRODUCTION

Chloroplast ribosomes originated from an ancestral cyanobacterium are the crossroad of RNA and protein metabolisms, essential for plant development and viability (1). Like its ancestor, chloroplast translation occurs in the 70S ribosomes comprising a large (50S) and a small (30S) ribosomal subunit (2,3). These subunits consist of multiple ribosomal proteins and a few ribosomal RNAs (rRNA). The chloroplast genome encodes half of the 30S ribosomal proteins and a quarter of the 50S ribosomal proteins (4). While most ribosomal proteins are indispensable for chloroplast biogenesis, some of them implicated in ribosome assembly and regulation (or optimization) of translation are nonessential (5,6). Despite its small number, rRNAs play the most crucial role in ribosome biogenesis (7). In fact, the crystal structure of 70S ribosome reveals that the catalytic activity seems to depend entirely on rRNA (8).

In bacteria, 16S and 23S rRNA are co-transcribed into polycistronic precursor and serve as a scaffold for ribosomal protein binding during the assembly of 30S and 50S ribosomal subunits, respectively (9). The rRNAs participate in controlling the fidelity of the codon-anticodon base pairing in the decoding center (16S rRNA in the 30S) and the catalytic peptidyl transferase activity (23S rRNA in the 50S) (9). Given the similarities of sequence and secondary structure of rRNAs in chloroplast and bacteria (10–12), chloroplast rRNAs are thought to have comparable characteristics with bacterial rRNAs. Like bacterial rRNA, the plastid-encoded bacterial-type RNA polymerase (PEP) produces polycistronic primary rRNAs, which further undergo maturation through multiple post-transcriptional processing, including cleavage, stabilization, and splicing (13–15).

Ribosome assembly is a highly coordinated process incorporating rRNA processing, nucleotide modification, addition and modification of ribosomal proteins, and binding and release of assembly factors (16). Many steps co-occur during or immediately after rRNA transcription (9,17). In *Escherichia coli*, the 30S subunit biogenesis requires parallel and hierarchical assembly pathways. Twenty ribosomal proteins (divided into primary-, secondary-, and tertiary-binding proteins) can be successively loaded to 16S rRNA or form the assembly intermediates in parallel to assemble a 30S ribosomal subunit (18). Assembly cofactors play a critical role in rRNA folding and protein-rRNA interactions, since a long rRNA can spontaneously form misfolded secondary structures (9). However, little is known about the assembly cofactors involved in the chloroplast 30S subunit. Among those characterized assembly cofactors, RBF1, a homolog of the bacterial RbfA (Ribosome-binding factor A), participates in the biogenesis of 30S subunits in chloroplasts by acting in both the 5’ and 3’ end processing of 16S rRNA (19). Another assembly cofactor, Embryo defective 15 (EMB15), whose N-terminal part contains the bacterial RimM (Ribosome maturation factor M)-like domain, has recently been discovered in maize (*Zea mays*) plastid. Loss of EMB15 severely impairs 16S rRNA maturation and retards embryo development at an early stage (20).

Methylation is the major post-transcriptional rRNA modification in all living organisms. In bacteria, ten methyltransferases involve in 16S rRNA methylation during ribosome assembly, which contributes to the fidelity of translation in most cases (21). Only three 16S rRNA methyltransferases resembling those in bacteria have been identified in the chloroplast (12,22,23). PALEFACE1 (PFC1), a homolog of bacterial ribosomal RNA small subunit methyltransferase A (RsmA), was identified in a cold-sensitive Arabidopsis mutant and is involved in the dimethylation of two adjacent adenines at the 3’ end of 16S rRNA (22). Likewise, the *E. coli* mutant lacking RsmA exhibits chilling-sensitive growth defects (24). Like *E.coli* RsmD (25), Arabidopsis RsmD (AtRsmD) methyltransferase catalyzes the *N^2^*-methylation of guanine 915 (m^2^G915) of 16S rRNA, an equivalent of m^2^G966 in *E. coli* 16S rRNA, and participates in the adaption to cold stress (12). Similar cold-sensitivity was reported in the *E.coli rsmD* mutant (26). Another chloroplast MraW-like protein (CMAL) catalyzes the *N*^4^-methylation of cytosine 1352 (m^4^C1402) of 16S rRNA in Arabidopsis (23), equivalent to the m^4^C1402 by *E. coli* MraW/RsmH (27). Loss of CMAL impairs 30S ribosome biogenesis, thus constraining plant development. The decreased auxin synthesis/signaling and photosynthesis rate are suggested as potential causes of the growth retardation, indicating that the reduced chloroplast translation affects other metabolic activity. Notably, the *E. coli rsmH* mutant shows only a mild growth deficiency compared to the *cmal* mutant (28), indicating an evolutionarily acquired neo-function of CMAL in plants. All these findings suggest that chloroplast 16S rRNA methylation is vital for translation, ribosome biogenesis, plant development, and acclimation under fluctuating environment conditions (29,30).

Ribosome assembly, rRNA processing, and protein translation are interdependent events, causing a chicken-and-egg problem, especially for conferring precise functionality of related proteins. In many cases, the disrupted rRNA processing results from the secondary effects (5,31). Nonetheless, the forward genetic approach can be one of the toolboxes identifying proteins involved in these interdependent processes. Interestingly, studies on the Arabidopsis *yellow variegated 2* (*var2*) mutant have unveiled various chloroplast components involved in protein biosynthesis (31–35). The *var2* mutant shows a defect in photosystem II (PSII) repair and chloroplast biogenesis owing to the lack of FtsH2 metalloprotease (36–39). Unlike other chloroplast protease mutants generally showing growth retardation, *var2* mutant develops variegated leaves from the emerging true leaf, whereas cotyledons remain green. To date, a dozen suppressors of *var2* (*svr*) have been reported, rescuing the leaf variegation. The identified genetic factors are primarily involved in chloroplast transcription (31) and translation (32,40), hence proving a so-called threshold model: reducing substrates of FtsH2 (e.g., PSII core proteins) results in less requirement of FtsH2, thereby restoring leaf greening (41). Given that rRNA processing and translation are coupled events, we considered that studies on *svr* mutants might shed new light on chloroplast ribosome biogenesis.

The present study identified a causal mutation in *svr12* through whole genome sequencing and complementation analyses. *SVR12* encodes the chloroplast-specific AtRsmD methyltransferase. Loss of AtRsmD explicitly compromises 16S rRNA maturation, but not 23 rRNA, and translation efficiency. The ensuing co-immunoprecipitation coupled with mass-spectrometry (CoIP-MS) analysis revealed the association of AtRsmD with multiple ribosomal proteins and maturation factors such as ribosome maturation factor M (AtRimM). Like the *atrsmD* mutant, loss of AtRimM substantially impaired 16S rRNA processing. Interestingly, the m^2^G915 level was dramatically decreased in the *atrimM* mutant, suggesting that AtRimM facilitates the m^2^G915 modification by interacting with AtRsmD. Here, we report a new post-transcriptional regulation of 16S rRNA maturation mediated by AtRsmD and AtRimM, a 16S rRNA methyltransferase and assembly cofactor.

## MATERIALS AND METHODS

### Plant materials and growth conditions

All *Arabidopsis* seeds used in this study were in the Columbia-0 (Col-0) ecotype and harvested from plants grown on soil under continuous light (CL, 100 μmol·m^-2^·s^−1^ of light from Cool White Fluorescent bulbs) at 22 ± 2°C. Arabidopsis T-DNA insertional knockout mutant seeds of *atrsmD* (SAIL_723_B06), *atrimM* (SALK_119256), and *var2-ko* (SAIL_253_A03) were obtained from the Nottingham Arabidopsis Stock Centre (NASC). Arabidopsis seed of the *var2-9* allele used in this study was reported previously (39,42). To generate double mutants of *var2-9 atrsmD, var2-ko atrsmD*, and *var2-9 atrimM*, the *atrsmD* or *atrimM* mutant was crossed with either *var2-9* or *var2-ko.* Homozygous plants were isolated from the segregating F_2_ plants by performing PCR-based genotyping with appropriate primers (Supplementary Table S1). Seeds were surface-sterilized with 1.6% (v/v) hypochlorite solution and plated on Murashige and Skoog (MS) medium (Duchefa Biochemie) containing 0.7% (w/v) Plant agar (Duchefa Biochemie). After a three-day stratification at 4°C in darkness, the seeds were placed in a growth chamber (CU-41L4; Percival Scientific) under CL (40 μmol·m^-2^·s^−1^ at 22°C).

### EMS mutagenesis and suppressor screen for *var2-9*

*var2-9* seeds were submerged in 0.4% (v/v) ethyl methanesulphonate (EMS; Sigma-Aldrich) solution for 8 hours with gentle agitation, washed several times in sterile water to remove the remaining EMS, and dried on Whatman filter papers. Approximately 10,000 M_1_ plants sub-grouped into forty were grown on soil under CL conditions, and M_2_ seeds were harvested as pools. Around 80,000 M_2_ plants were then germinated on soil, mutant plants showing an entirely recovered leaf variegation phenotype were selected, and their seeds were individually harvested. More than a hundred of independent *var2-9* suppressor (*svr*) lines were initially selected. Among them, *svr12* was chosen and genetically crossed with parental *var2-9* plants. The suppressor phenotype of the crossed line showed a 1:3 segregation in the F_2_ generation, indicating a single recessive trait.

### Mapping of *svr12* by whole-genome sequencing

To establish a mapping population, *svr12* was once backcrossed to the *var2-9* parental line. Fifty F_2_ plants that displayed the suppressor phenotype were chosen and pooled. Genomic DNAs from the pool and *var2-9* were extracted with the DNeasy Plant kit (Qiagen). One μg of genomic DNA was used for library construction with the Truseq DNA PCR-free kit (Illumina) following the manufacturer’s instructions for 350 bp insert size. Next, paired-end sequencing was performed on a HiSeq 2500 sequencer (Illumina), yielding 125 bp paired-end reads. Genome resequencing data were first processed by SolexaQA (43) and Cutadapt to remove low-quality regions and adapter sequences, respectively. Clean reads were then mapped to the TAIR10 genome by BWA-MEM (44) with default parameters. Single nucleotide polymorphisms (SNPs) were called by producing mpileup files with SAMtools (45), and poor quality SNPs with a mapping quality less than 60 or depth less than 3 or over 200 were filtered out by VCFtools (46). SNPs found only in the *svr12* mutant pool were retained with the assumption that these mutations were generated by EMS mutagenesis. SHOREmap 3.0 (47) was used to draw the allele frequency graph and identify the region that contained the causal mutation surrounded by tightly linked SNPs. In the identified region, genes containing at least one novel mutation were regarded as candidate genes.

### Generating Arabidopsis transgenic plants

The stop-codon-less *AtRsmD* coding sequence (CDS) and the stop-codon-less genomic *AtRsmD* DNA containing the 1-kb promoter region were cloned into a pDONR/Zeo entry vector (Thermo Fisher Scientific) via Gateway BP reaction (Thermo Fisher Scientific) and subsequently recombined through the Gateway LR reaction (Thermo Fisher Scientific) into the destination vectors pGWB505 (48) for C-terminal fusion with GFP and pGWB516 (48) for C-terminal fusion with 4×Myc to create *p35S:AtRsmD-GFP* and *pAtRsmD:AtRsmD-4×Myc* constructs, respectively. The generated vectors were transformed into *Agrobacterium tumefaciens* strain GV3101 using a MicroPulser Electroporation system (Bio-Rad). After delivering the generated vectors to *svr12* or *atrmsD* mutant plants using *Agrobacterium-mediated* transformation by the floral dip method (49), homozygous transgenic plants were selected on MS medium containing 35 mg/L hygromycin (Thermo Fisher Scientific).

### Chloroplast purification and subfractionation

Intact chloroplasts were isolated from the foliar tissues of 3-week-old *atrsmD* transgenic plants expressing *pAtRsmD:AtRsmD-4×Myc* grown under CL conditions as described previously (38). Isolated chloroplasts were subfractionated into stroma, envelope membrane, and thylakoid fractions according to the method described previously (50,51). Briefly, purified intact chloroplasts were incubated in hypotonic lysis buffer (25 mM HEPES-KOH [pH 8.0) containing 1× Complete protease inhibitor cocktail [Roche]) for 1 h at 4°C with gentle agitation. The lysate was loaded on top of a 25 mM HEPES-buffered sucrose step gradient (3.6 mL, 1.2 M; 3.6 mL, 1.0 M; and 3.6 mL, 0.46 M, respectively) in a 12.5 mL ultracentrifuge tube and centrifuged for 1 h at 4°C at 200,000 × *g* in a swinging bucket rotor (Sorval SW-41 Ti; Beckman Coulter), with moderate acceleration and deceleration. The soluble stroma fraction was collected from the top of the gradient. The envelope membrane and thylakoid fractions were obtained from the 0.46 M/1.0 M sucrose interface and 1.0 M/1.2 M sucrose interface, respectively. After precipitating chloroplasts and stroma subfractions using the methanol/chloroform method, the resulting pellets were resuspended in equal volumes of SDS sample buffer.

### Confocal laser scanning microscopy

To monitor subcellular localization of the AtRsmD protein, cotyledons from 5-day-old transgenic *svr12* seedlings over-expressing the *p35S:AtRsmD-GFP* were used for confocal laser scanning microscopy (CLSM) analysis using a TCS SP8 (Leica Microsystems). GFP fluorescence was excited with an argon laser at a wavelength of 488 nm, and the emission of GFP fluorescence was collected at between 500 and 530 nm. Chlorophyll autofluorescence was obtained between 650 and 700 nm. Processed images were obtained using LAS AF Lite software, version 2.6.3 (Leica Microsystems).

### RT-qPCR analysis

Total RNA (1 μg) extracted from three independent biological replicates of the indicated genotypes using the Universal Plant Total RNA Extraction Kit (Spin-column) (BioTeke) was reverse-transcribed with the HiScript II Q RT SuperMix (Vazyme Biotech) according to the manufacturer’s recommendations. The RT-qPCR was performed using the ChamQ Universal qPCR Master Mix (Vazyme Biotech) on the QuantStudio™ 6 Flex Real-Time PCR System (Applied Biosystems). Relative transcript abundances of the selected genes were calculated with the ddCt method (52) and normalized to those of the *PP2A* (At1g13320) gene. The primer sequences used for RT-qPCR are listed in Supplementary Table S1.

### RNA immunoprecipitation analysis

Endogenous RNAs were co-immunoprecipitated with AtRsmD-GFP proteins, as described previously (53,54), with minor modifications. Briefly, RNA-protein complexes in approximately 1 g fresh weight of 5-day-old seedlings of *svr12 p35S:AtRsmD-GFP* and wild-type *p35S:GFP* were cross-linked by 1% (v/v) formaldehyde for 15 min and quenched with 125 mM glycine for 15 min. The seedlings were washed thrice with RNase-free water at 4°C, frozen in liquid nitrogen, ground to fine powder, and homogenized in lysis buffer containing 20 mM Tris-HCl (pH 7.5), 150 mM NaCl, 1 mM MgCl_2_, 1 mM CaCl_2_, 0.1% (w/v) SDS, 1% (w/v) sodium deoxycholate, 1% (v/v) Triton X-100, 1 mM PMSF, 5 mM DTT, and 1× Complete protease inhibitor cocktail (Roche). After centrifuging at 16,000 × g for 7 min at 4°C, the supernatants diluted 10 times with dilution buffer (10 mM Tris-HCl [pH 7.5], 150 mM NaCl, 0.5 mM EDTA) were incubated with 20 μL of GFP-Trap magnetic agarose beads (GFP-TrapMA; Chromotek) for 12 h at 4°C by vertical rotation. The beads were washed 5 times with washing buffer (50 mM Tris-HCl [pH 7.5], 500 mM NaCl, 4 mM MgCl_2_, 0.5% [w/v] sodium deoxycholate, 0.1% [w/v] SDS, 2 M Urea, 2 mM DTT) at 4°C. RNAs were eluted from the beads with TRI-Reagent™ solution (Thermo Fisher Scientific), precipitated by adding 1 volume of isopropanol and 0.1 volume of 3 M sodium acetate (pH 5.2) for 10 min at −20°C, and washed with 75% (v/v) ethanol. The resulting RNA pellets were dissolved in 12 μL of RNase-free water. For the input samples, RNAs were also extracted from the cell lysates following the same procedure. RT-qPCR was then performed with immunoprecipitated and input RNAs. The primer sequences of chloroplast 16S rRNA and *psbA* (used as a negative control) are listed in Supplementary Table S1. The amount of RNA co-precipitated with AtRsmD-GFP was calculated in comparison with the respective input RNA used for each RNA immunoprecipitation (RIP), as described previously (54).

### Primer extension assay

Five hundred ng of total RNA, extracted from the indicated genotypes, as described above, was used to perform the primer extension assay. Briefly, the gene-specific primer was designed with the reverse-complementary sequence for the target region (between 963 and 979; 5’-AAGGCACCCCTCTCTTT-3’) of the chloroplast 16S rRNA. The primer and DNA marker were end-labeled with 10 μCi of [γ-^32^P]ATP by T4 polynucleotide kinase (Promega) and then used for reverse transcription with avian myeloblastosis virus (AMV) reverse transcriptase (Promega) according to the manufacturer’s instructions. The products of reverse transcription were separated on 8% denaturing polyacrylamide gel containing 40% polyacrylamide/bis (19:1) solution, 7M urea, and 1× TBE buffer. The gel was washed with a fixation solution [1× TBE buffer, 10% (v/v) ethanol, 10% (v/v) methanol], and then the radioactivity was visualized with a Personal Molecular Imager (PMI) system (Bio-Rad). To quantify the levels of endogenous m^2^G915 in chloroplast 16S rRNA, primer extension products reverse-transcribed with a gene-specific primer (reverse-complementary to the 16S rRNA nucleotides 1092-1108; 5’-CAGTCTGTTCAGGGTTC-3’) and AMV reverse transcriptase (Promega) were analyzed by qPCR with the primer pairs, as described in Supplementary Figure S4B, and the primer sequences are listed in Supplementary Table S1.

### Immunoblot analysis

Immunoblot analysis was performed as described before (39). Briefly, total protein extracted from seedlings grown on MS agar medium with homogenization buffer (0.0625 M Tris-HCl [pH 6.8], 10% [v/v] glycerol, 1% [w/v] SDS, 0.01% [v/v] β-mercaptoethanol) was separated by 10% SDS-PAGE gel and blotted onto Immun-Blot PVDF membrane (Bio-Rad). D1, D2, CP43, RbcL, RPS1, PRL2, RPL4, and LHCB4 proteins were immunochemically detected with rabbit anti-D1 (1:10,000; Agrisera), rabbit anti-D2 (1:10,000; Agrisera), rabbit anti-CP43 (1:10,000; Agrisera), rabbit anti-RbcL (1:10,000; Agrisera), rabbit anti-RPS1 (1:10,000; PhytoAB), rabbit anti-RPL2 (1:10,000; PhytoAB), rabbit anti-PRL4 (1:10,000; PhytoAB), and rabbit anti-LHCB4 (1:7,000; Agrisera) polyclonal antibodies, respectively. AtRsmD-4×Myc fusion proteins were detected using a mouse anti-Myc monoclonal antibody (1:10,000; Cell Signaling Technology).

### CoIP, BiFC, and MS analyses

To identify proteins associated with AtRsmD, co-immunoprecipitation coupled with mass-spectrometry (CoIP-MS) analysis was performed as described previously (55) with minor modifications. Briefly, 5-day-old *svr12* and WT transgenic seedlings harboring the *p35S:AtRsmD-GFP* were homogenized to fine powder in liquid nitrogen with mortar and pestle. Approximately 1 g of powder was used to extract total proteins with IP buffer containing 20 mM HEPES-KOH (pH 7.4), 2 mM EDTA, 2 mM EGTA, 25 mM NaF, 1 mM NA3VO4, 10% (v/v) glycerol, 0.1 M NaCl, 0.5% (v/v) Triton X-100, and 1× Complete protease inhibitor cocktail (Roche). After protein extraction, 10 mg of the total proteins were incubated with 20 μL of GFP-Trap magnetic agarose beads (GFP-TrapMA; Chromotek) for 2 hours at 4°C by vertical rotation. The beads were then washed four times with washing buffer (IP buffer without Triton X-100) and washed twice with 1× PBS buffer. Afterward, the beads were subjected to direct on-bead trypsin digestion followed by MS analysis. The same CoIP-MS analysis was also performed with total protein extracted from wild-type transgenic plants overexpressing free GFP for negative control.

CoIP and BiFC assays were carried out with *Nicotiana benthamiana* leaves as described (56). Briefly, two chloroplast proteins, AtRimM and RPS20, were selected from the 69 putative AtRsmD-associated proteins (Supplementary Table S2) identified through CoIP-MS analyses in *svr12 p35S:AtRsmD-GFP* plants but absent in the GFP control. RbcS1A, a member of the Rubisco small subunit, was used as the negative control. For BiFC assays, the pDONR/Zeo entry vectors containing the stop-codon-less CDSs of *AtRsmD, AtRimM, RPS20*, and *RbcS1A* were recombined into the split-YFP vectors (pGTQL1211 or 1221) (57) through Gateway LR reaction (Thermo Fisher Scientific). After the infiltration of *A. tumefaciens* mixtures harboring the appropriate BiFC constructs with a 1 mL syringe into the abaxial side of 4-week-old *N. benthamiana* leaves, the YFP fluorescent signals were monitored by TCS SP8 (Leica Microsystems) at an excitation wavelength of 514 nm. For CoIP assays, the same entry vectors used for BiFC were recombined into the destination vector pGWB617 for C-terminal fusion with 4×Myc. Each construct was transformed into *A. tumefaciens* and transiently coexpressed with *p35S:AtRsmD-GFP* construct in 4-week-old *N. benthamiana* leaves. After the CoIP assays, as described above, the beads were suspended with 2× SDS sample buffer and incubated at 95°C for 10 min. The eluates were subjected to 10% SDS-PAGE gels, and the interaction between AtRsmD and the selected proteins was determined by immunoblot analysis with a mouse anti-GFP monoclonal antibody (1:5,000; Roche) and a mouse anti-Myc monoclonal antibody (1:10,000; Cell Signaling Technology), respectively.

### RNA gel blot and polysome analyses

One μg of total RNAs extracted from 5-day-old seedlings using TRI-Reagent^TM^ solution (Thermo Fisher Scientific) was subjected to RNA gel blot analyses. Briefly, each RNA solution was mixed with 2× RNA loading dye (Thermo Fisher Scientific), heated for 10 min at 70°C, and immediately placed on ice. After loading the mixtures onto 1% formaldehyde agarose gels, the gels were run in 1× RNA running buffer (1 × MOPS and 6% [v/v] formaldehyde) and blotted onto the Hybond-N^+^ membrane (GE Healthcare) soaked in 20× SSC (3.0 M NaCl and 0.3 sodium citrate). After UV cross-linking using a Stratalinker 2400 UV Crosslinker (1,000 microjoules × 100 for 50s], the membranes were hybridized in 10 mL hybridization buffer (Sigma-Aldrich) containing 1 μM of biotin-conjugated RNA probes for 4 h at 42°C. The 3’-biotin-labeled RNA probes were synthesized and purified by high pressure liquid chromatography (HPLC) (Jie Li Biology, China), and the probe sequences are listed in Supplementary Table S1. The hybridized membranes were washed twice for 5 min at 42°C with low stringency wash buffer (2× SSC, 0.1% [w/v] SDS), and subsequently washed twice for 10 min at 42°C with ultra-high wash buffer (0.1× SSC, 0.1% [w/v] SDS). The biotin-labeled probes hybridized to RNAs were detected using a Chemiluminescent Nucleic Acid Detection Module kit (Thermo Fisher Scientific) according to the manufacturer’s instructions.

Polysome analyses were conducted using 5-day-old seedlings as previously described (58), with minor modifications. Briefly, 0.3 mg of cotyledons were frozen in liquid nitrogen, ground to fine powder, and homogenized in 1 mL polysome extraction buffer containing 0.2 M Tris-HCl (pH 9.0), 0.2 M KCl, 35 mM MgCl_2_, 25 mM EGTA, 0.2 M sucrose, 1% (v/v) Triton X-100, 2% (v/v) polyoxyethylene-10-tridecyl ether, heparin 0.5 mg/mL, 100 mM β-mercaptoethanol, 100 μg/mL chloramphenicol, and 25 μg/mL cycloheximide. The lysates were passed through 100 μm cell strainer (Falcon) and centrifuged at 850× g for 5 min at 4°C to eliminate debris. After adding 0.5% (w/v) sodium deoxycholate to the supernatants, the remaining insoluble material was removed by centrifugation at 16,000× g for 15 min at 4°C. The supernatant (0.6 mL) was gently layered on top of 4.4 mL sucrose gradient solution (15 to 55%) in a 5.1 mL Quick-Seal polypropylene tube (Beckman Coulter) and centrifuged at 45,000 rpm for 65 min at 4°C (Sorval NVT 90; Beckman Coulter). In parallel, the lysates supplemented with 20mM EDTA were also subjected to sucrose gradient solution containing 0.1 mM EDTA instead of MgCl_2_. RNA was extracted from 200 μL of each sucrose gradient fraction with the TRI-Reagent^TM^ solution. Each fraction was mixed with the 4× SDS sample buffer for protein isolation.

### Determining photochemical efficiency and chlorophyll content

Photochemical efficiency of PSII (Fv/Fm) was determined using a FluorCam system (FC800-C/1010GFP; Photon Systems Instruments) equipped with a CCD camera and an irradiation system according to the instrument manufacturer’s instructions. Total chlorophyll was extracted from leaves by boiling them in 95% (v/v) ethanol at 80°C for 20 minutes. The relative chlorophyll content per fresh weight was calculated as described by Lichtenthaler (59).

### Homology modeling

The three-dimensional (3D) structure of AdoMet-MTase domain (amino acid sequence 114-303) of AtRsmD classified by NCBI (domain architecture ID 10508003) was modeled using the SWISS MODEL (60) and PyMOL software, version 2.0 (The PyMOL Molecular Graphics System). The modeling template was obtained from the experimentally-determined 3D structure of *E. coli* RsmD (amino acids 12-187; PBD:2FPO) (25).

## RESULTS

### Loss of AtRsmD rescues the leaf variegation in *var2*

To explore chloroplast components involved in ribosome biogenesis and proteostasis, we mutagenized *var2-9* (one of the *var2* alleles) harboring a missense mutation in FtsH2 that impairs its substrate-unfolding activity (39). We isolated several suppressors in the M_2_ population of ethyl methanesulfonate (EMS)-mutagenized *var2-9* (Figure 1A). Among them, *svr12* exhibited a leafy phenotype with the restored maximum photochemical efficiency of PSII (Fv/Fm) and total chlorophyll content (Figure 1B and C). Whole-genome sequencing analyses revealed that the *svr12* genome contains a single nucleotide polymorphism with a G-to-A mutation in a gene encoding the chloroplast AtRsmD methyltransferase that results in a nonsense mutation at codon 30, Glutamine(Q)30 to stop codon (Supplementary Figure S1). The subsequent complementation assay by expressing Myc-tagged AtRsmD under the control of its native promoter (*pAtRsmD:AtRsmD-Myc*) confirmed the causal relation between the nonsense mutation and the suppression of *var2-9* phenotypes (Figure 1D to 1F). Loss of AtRsmD also fully rescued the foliar phenotype of *var2-9* and *var2* null (*var2-ko*) alleles (Figure 1G), indicating that AtRsmD is indispensable for *var2* phenotypes.

**Figure 1.**
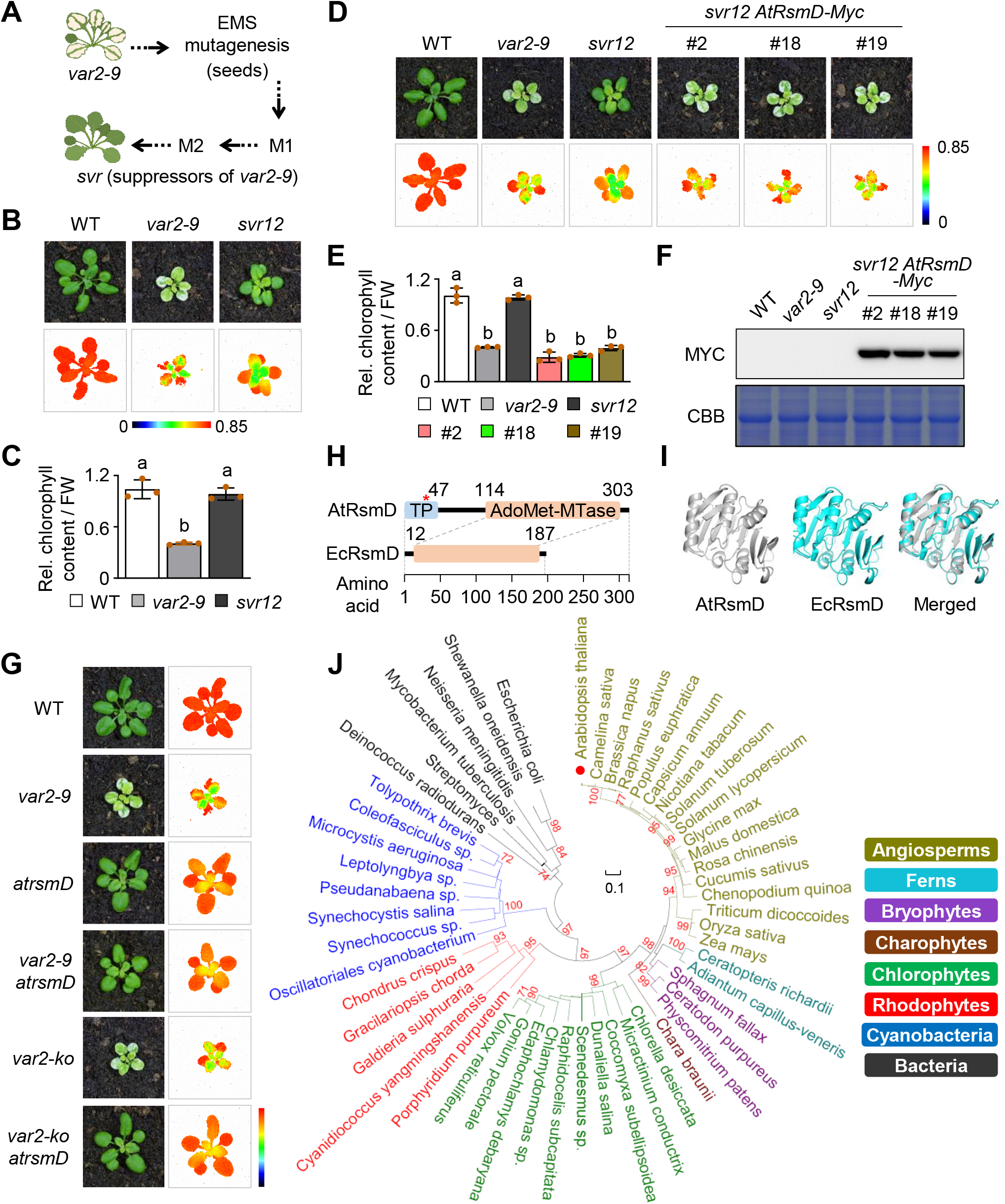
Loss of chloroplast AtRsmD suppresses leaf variegation in *var2* mutants. (**A**) Schematic illustration of the genetic screening for *var2* suppressors. Around 80,000 EMS-mutagenized M_2_ plants derived from approximately 10,000 M_1_ plants were screened to isolate suppressor lines (more details in MATERIALS AND METHODS). (**B**) and (**C**) Foliar phenotype (**B**, top panels), maximum photochemical efficiency of PSII (Fv/Fm; **B**, bottom panels), and total chlorophyll content (**C**) were examined in 3-week-old wild-type (WT), *var2-9*, and *svr12* plants grown on soil under continuous light (CL). More than 20 plants were used for each experiment, and the representative images are shown at the same scale. (**D**) to (**F**) Complementation assay by expressing Myc-tagged AtRsmD under the control of its native promoter (*pAtRsmD:AtRsmD-Myc*) in *svr12.* Foliar phenotype (**D**, top panels), Fv/Fm (**D**, bottom panels), total chlorophyll content (**E**), and protein abundance of AtRsmD-Myc (**F**) were examined in 3-week-old plants of WT, *var2-9, svr12*, and three independent *svr12 pAtRsmD:AtRsmD-Myc* (*svr12 AtRsmD-Myc*) lines grown under CL. Myc antibody was used for the immunoblot analysis, and SDS-PAGE gel stained with coomassie brilliant blue (CBB) was used as a loading control (**F**). In (**C**) and (**E**), data are presented as mean ± SD (*n*=3), and lowercase letters indicate statistically significant differences between mean values (*P* < 0.01, one-way ANOVA with posthoc Tukey’s HSD test). (**G**) Foliar phenotype (left panels) and Fv/Fm (right panels) were examined in CL-grown 3-week-old WT, *var2-9, var2-ko, atrsmD, var2-9 atrsmD*, and *var2-ko atrsmD.* (**H**) Diagram shows AtRsmD containing an N-terminal chloroplast transit peptide (TP; amino acids 1-47) and a C-terminal AdoMet-MTase domain (amino acids 114-303) similar to *E. coli* RsmD (EcRsmD; amino acids 12-187). The red asterisk on the TP indicates the nonsense mutation (Q30 to stop codon) found in *svr12.* (**I**) Three-dimensional (3D) structures of AdoMet-MTase domain of AtRsmD and EcRsmD modeled by the PyMoL software. Experimentally determined 3D structure of EcRsmD (PBD: 2FPO) was used as a template for the modeling. (**J**) Phylogenetic analysis of AtRsmD orthologs from different species, including representative bacteria, cyanobacteria, rhodophytes, chlorophytes, charophytes, bryophytes, ferns, and angiosperms as indicated by different color codes. The phylogenetic tree was made using the MEGA8 software, and the scale bar represents 0.1 estimated amino acid substitutions per site.

The AtRsmD protein contains a predictable chloroplast transit peptide at the N-terminus and a C-terminal region resembling S-adenosylmethionine-dependent methyltransferase (AdoMet-MTase) in *E. coli* (Figure 1H). Homology modeling showed identical structures of the AdoMet-MTase domain in AtRsmD and *E.coli* RsmD (Figure 1I). Multiple sequence alignment revealed that the AdoMet-MTase domain of AtRsmD is highly conserved across different bacterial species, including cyanobacterium *Synechococcus* (Supplementary Figure S2). Intriguingly, the phylogenetic analysis showed that in eukaryotes, AtRsmD orthologs are only present without gene duplication in photosynthetic organisms, including plants, mosses, and red/green algae, but not in yeast or animals (Figure 1J). This suggests a conserved chloroplast-specific functionality of RsmD during the evolution of Viridiplantae. Consistent with the phylogenetic data, we found that the CaMV 35S promoter-driven GFP (Green Fluorescent Protein)-tagged AtRsmD only appeared in chloroplasts (Supplementary Figure S3A), which confirms the previous observation (12). In addition, the chloroplast subfractionation assay using Arabidopsis *atrsmD* transgenic plants expressing *pAtRsmD:AtRsmD-Myc* revealed its exclusive localization in the stroma as evidenced by the similar chloroplastic distribution of the stroma marker protein Rubisco large subunit (RbcL) (Supplementary Figure S3B).

### AtRsmD binds the chloroplast *16S* rRNA for m^2^G915 modification

*E. coli* RsmD is an AdoMet-MTase catalyzing the methylation of G966 to m^2^G966 in *16S* rRNA (25,61). Since the RsmD orthologs were found in most photosynthetic organisms (Figure 1J), and the sequence and secondary structure of chloroplast 16S rRNA are highly comparable to those of *E. coli 16S* rRNA (10–12), we assumed that AtRsmD would also methylate 16S rRNA in the chloroplast. By using a primer extension assay, a recent study reported that AtRsmD is indeed required for 16S rRNA m^2^G915 modification (corresponding to G966 of the *E. coli* 16S rRNA) (12). To quantify m^2^G915 modification, we first validated AtRsmD-16S rRNA interaction by RNA-immunoprecipitation coupled with qPCR (RIP-qPCR) analyses in *svr12 p35S:AtRsmD-GFP* transgenic plants. We included a wild-type (WT) transgenic plant expressing free GFP driven by the 35S promoter (p35S:GFP) as a negative control. The resulting RIP-qPCR assay confirmed the AtRsmD-16S rRNA interaction (Supplementary Figure S4A). Next, to investigate the catalytic activity of AtRsmD towards 16S rRNA m^2^G915 modification, we performed a primer extension assay using total RNAs isolated from seedlings of WT, *atrsmD*, and *atrsmD pAtRsmD:AtRsmD-Myc.* The primer extension might stop when the reverse transcriptase reaches the nucleotide preceding m^2^G915 because the methyl group blocks any G-C Watson-Crick base pairing (Supplementary Figure S4B). Accordingly, primer extension by reverse transcriptase stopped at C916 of WT 16S rRNA but not of *atrsmD* mutant 16S rRNA, which became significantly reverted by complementation with AtRsmD-Myc (Supplementary Figure S4C). This result indicates that AtRsmD is critical to methylate G915 chloroplast 16S rRNA, consistent with the recent study (12). The quantitative analysis of m^2^G915 level via a primer extension coupled to qPCR (primer extension-qPCR) assay (Supplementary Figure S4B) further confirmed that the transcript abundance of chloroplast 16S rRNA containing m^2^G915 was drastically decreased in *atrsmD* mutant compared to WT and *atrsmD pAtRsmD:AtRsmD-Myc* (Supplementary Figure S4D).

### AtRsmD is associated with the chloroplast 30S ribosomal subunit

Given that 16S rRNA is crucial in the assembly of the 30S ribosomal subunit in bacteria, we anticipated that AtRsmD might stably or transiently associate with the 30S ribosomal subunit to catalyze 16S rRNA m^2^G915 modification. To this end, we carried out co-immunoprecipitation (CoIP) coupled to mass spectrometry (MS) analyses to reveal its associated interactome. Two independent CoIP-MS analyses identified 69 proteins co-immunoprecipitated with the AtRsmD-GFP fusion protein but not with free GFP via a GFP-Trap coupled to magnetic agarose beads (Figure 2A; Supplementary Table S2). Like AtRsmD-Myc, also the AtRsmD-GFP fusion protein restored the leaf variegation phenotype in *svr12* (Supplementary Figure S5). Nonetheless, among the 69 identified proteins, only 28 proteins were predicted to localize to chloroplasts, 14 of them are either ribosomal proteins or proteins involved in 16S or 23S rRNA processing (Supplementary Table S2). The subsequent CoIP-immunoblot (Figure 2B) and bimolecular fluorescence complementation (BiFC; Figure 2C) analyses verified that AtRsmD interacts with at least two ribosomal factors, such as 30S ribosomal protein S20 (RPS20) and ribosome maturation factor M (AtRimM). By contrast, no interaction was observed with the Rubisco small subunit 1A (RbcS1A) protein (Figure 2B and C), corroborating the association of AtRsmD with the 30S ribosomal subunit.

**Figure 2.**
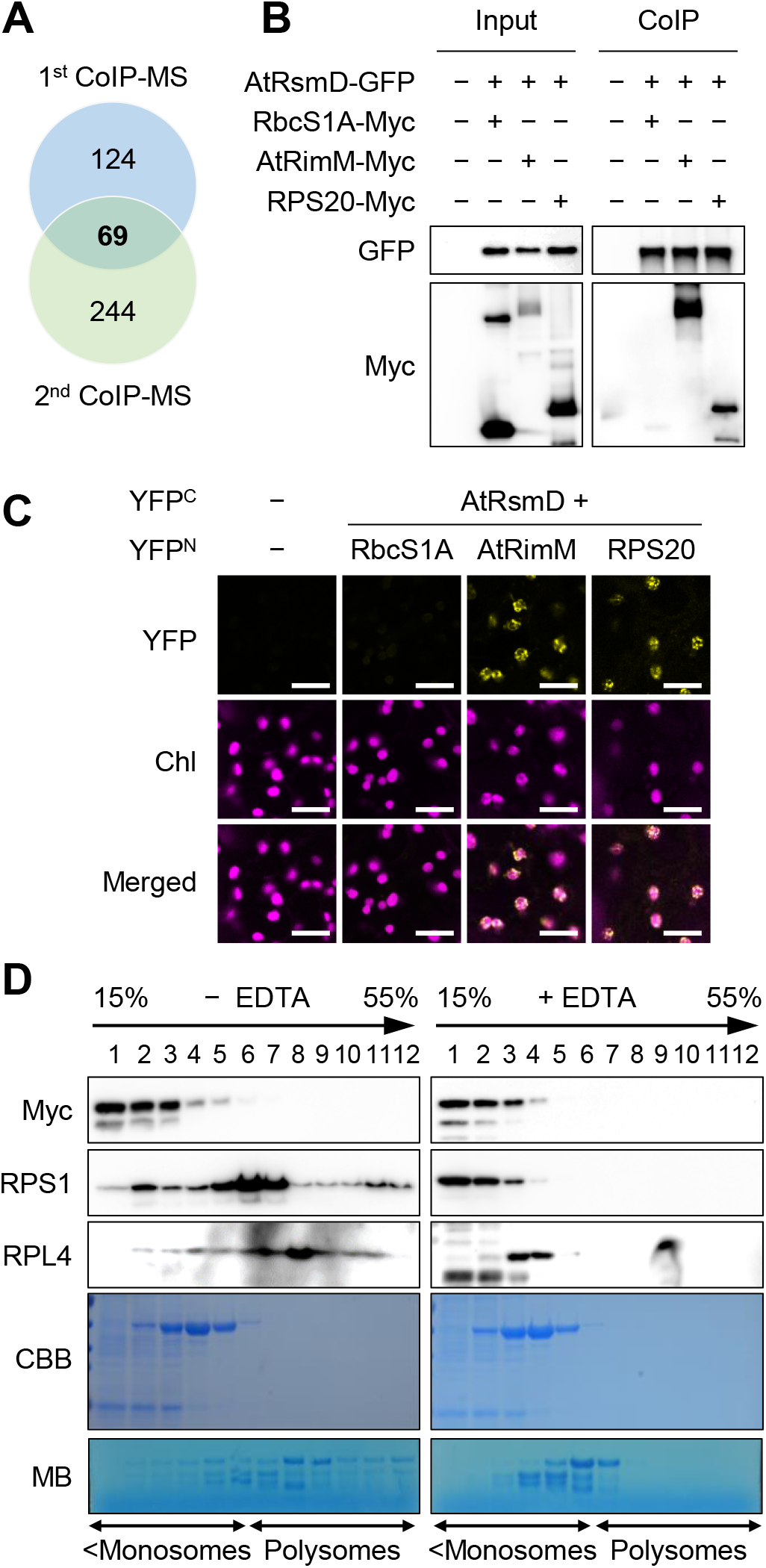
AtRsmD is associated with the chloroplast 30S ribosomal subunit and the ribosome maturation factor AtRimM. (**A**) Venn diagram showing the numbers of AtRsmD-associated proteins obtained from two independent CoIP-MS analyses. CL-grown 5-day-old seedlings of *svr12 p35S:AtRsmD-GFP* and WT *p35S:GFP* (negative control) were used. Only proteins coimmunoprecipitated with AtRsmD-GFP but not with free GFP were counted. (**B**) CoIP-immunoblot analyses with *N. benthamiana* leaves transiently coexpressing AtRsmD-GFP with RPS20-Myc or AtRimM-Myc. CoIP was performed with GFP-Trap beads, and the interaction was evaluated using the Myc antibody. (**C**) BiFC assays. YFP constructs of AtRsmD fused with the C-terminal part of YFP (YFPC) and RbcS1A, RPS20, and AtRimM fused with the N-terminal part of YFP (YFPN) were transiently coexpressed in *N. benthamiana* leaves in the indicated combinations. The fluorescence signal of the integrated YFP protein was observed by confocal microscopy. The images were taken at the same scale (scale bars: 20 μm). Chl, chlorophyll. RbcS1A was used for negative control for CoIP and BiFC assays. (**D**) AtRsmD association with free 30S ribosomal complex. Polysomes from 5-day-old CL-grown seedlings of *atrsmD pAtRsmD:AtRsmD-Myc* were separated by sucrose density gradient (15% to 55%) ultracentrifugation in the presence or absence of EDTA. The fractions were subjected to immunoblot analyses using antibodies against Myc and 30S (RPS1) and 50S (RPL4) ribosomal proteins and RNA gel blot analyses. Lanes 1 to 12 indicate the fractions on sucrose gradients from the top (15%) to the bottom (55%). The SDS-PAGE gels and RNA membranes were stained with CBB and methylene blue (MB), respectively. The sedimentation of the monosomes (fractions 1-6, containing immature ribosomal particles and monosomes) and polysomes (fractions 7-12) on the sucrose gradients was confirmed with the EDTA-treated control.

Besides, the sucrose density gradient fractionation coupled with immunoblot analyses found that AtRsmD is explicitly present in the low-sucrose density fraction (fractions 1-6), where immature ribosomal particles (such as free 30S and 50S subunits) and monosomes are sedimented. In contrast, the 30S ribosomal protein S1 (RPS1) and 50S ribosomal protein L4 (RPL4) also sedimented in fractions containing polysomes (fractions 7-12). Note that most AtRsmD-Myc were in fractions 1-3 with RPS1being present but almost no RPL4 (Figure 2D), suggesting that AtRsmD may primarily be associated with free 30S subunit but not with monosomes. Consistent with this observation, upon treating cell lysates with ribosome-dissociating reagent EDTA prior to the sucrose-density gradient fractionation, AtRsmD and RPS1 but not RPL4 were detected in the same fractions with the lowest sucrose concentrations (fractions 1-2). This result further affirms that AtRsmD is mainly associated with the free 30S ribosomal subunit.

### AtRimM is required for 16S rRNA m^2^G915 modification

Secondary structure of *E. coli* 16S rRNA shows that the RsmD target (G966) is located in helix 31, near the P-site, where the anticodon of tRNA with its amino acid pairs up with the corresponding mRNA codon (62,63). Helices 31 and 33b of 16S rRNA interact with RPS13 and RPS19, respectively, and RPS13 directly binds to RPS19, leading to the interaction with the anticodon stem-loops of P- and E-site tRNAs (64,65). In *E. coli*, RPS19 in the 30S subunit is essential for RsmD binding to the 16S rRNA (61). Given that AtRsmD appeared to be associated with RPS13 and AtRimM (Supplementary Table S2) and that RimM aids the association of RPS19 into the 30S ribosomal subunit during maturation in *E.coli* and maize (20,64,66), we presumed that AtRimM might also involve in the m^2^G915 modification by promoting AtRsmD binding to the 16S rRNA. Indeed, the primer extension-qPCR assay found that loss of AtRimM significantly reduces the m^2^G915 modification level in *atrimM* mutant versus WT seedlings (Figure 3), suggesting an additional function of AtRimM in facilitating the AtRsmD-mediated m^2^G915 modification.

**Figure 3.**
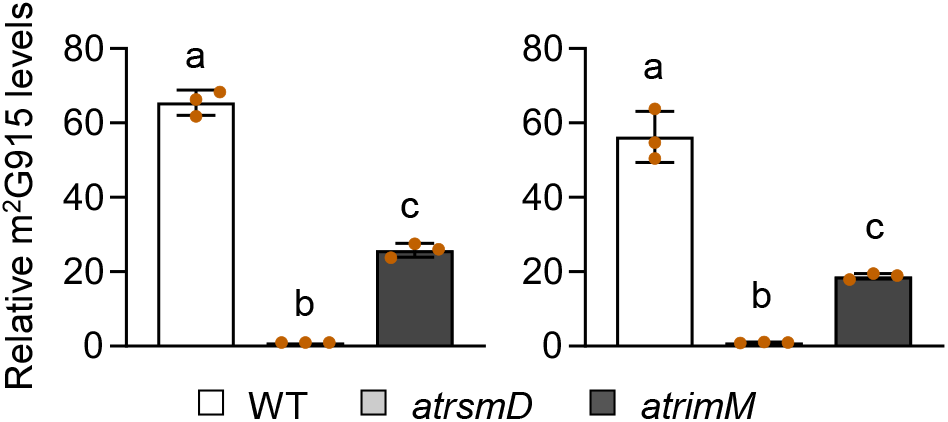
*AtRimM* mutation attenuates AtRsmD-dependent m^2^G915 modification. Primer extension coupled with qPCR analyses using total RNA extracted from CL-grown 5-day-old WT, *atrsmD*, and *atrimM* seedlings were performed as described in Supplementary Figure S4B. The experiment was repeated two times as shown, and data are presented as mean ± SD (*n*=3). Lowercase letters indicate statistically significant differences between mean values (*P* < 0.01, one-way ANOVA with posthoc Tukey’s HSD test).

### 16S rRNA m^2^G915 modification facilitates the assembly of 30S ribosomal subunit

The primary role of bacterial 16S rRNA methylations is to properly assemble ribosomes by contributing to the maturation and stabilization of 16S rRNA (7,67,68). Therefore, the interplay of AtRsmD and AtRimM towards m^2^G915 modification might be critical for the assembly and integrity of ribosomes. To this end, we examined the steady-state levels of chloroplast 30S and 50S ribosomal proteins in seedlings of WT, *atrsmD*, and *atrimM.* The result showed that the levels of 30S (RPS1) and 50S (RPL2 and RPL4) ribosomal proteins were significantly decreased in both *atrsmD* and *atrimM* relative to WT (Figure 4A).

**Figure 4.**
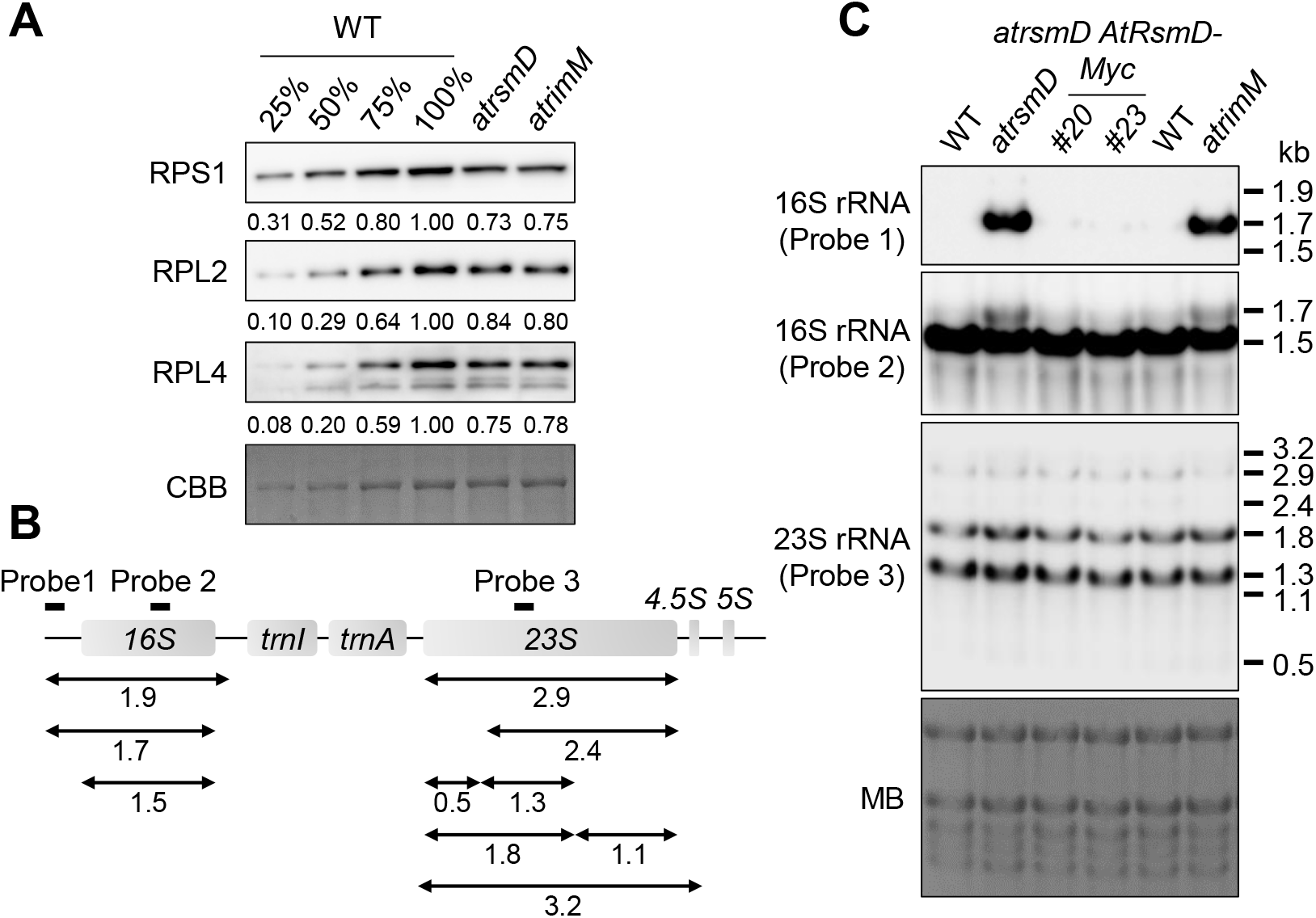
Ribosomal protein accumulation and 16S rRNA maturation are compromised in *atrsmD* and *atrimM* mutants. (**A**) Abundances of 30S (RPS1) and 50S (RPL1 and RPL4) ribosomal proteins were determined in 5-day-old seedlings of WT, *atrsmD*, and *atrimM* grown under CL conditions by immunoblot analyses. SDS-PAGE gel stained with CBB was used as the loading control. (**B**) Schematic diagram showing the chloroplast rRNA operon and the probe positions used for RNA gel blot analyses. Sizes of the major accumulating 16S and 23S rRNA transcripts are indicated in kilobase pairs (kb) below the map. (**C**) RNA gel blot analyses showing the processing patterns of chloroplast 16S and 23S rRNAs in CL-grown 5-day-old WT, *atrsmD*, and *atrimM* seedlings. RNA membrane stained with MB was used as a loading control.

Since 16S rRNA maturation is critical for ribosome assembly (14), we wondered whether the m^2^G915 modification can affect 16S rRNA metabolism. We first designed a set of probes to detect the precursor and mature forms of 16S and 23S rRNAs (Figure 4B). Then, RNA gel blot analyses were performed with total RNAs extracted from seedlings of WT, *atrsmD, atrsmD pAtRsmD:AtRsmD-Myc*, and *atrimM.* The results showed that the abundance of the 16S rRNA precursor (1.7 kb) was remarkably increased in both *atrsmD* and *atrimM* compared to WT and *atrsmD pAtRsmD:AtRsmD-Myc*, whereas the level of the processed 1.5 kb 16S rRNA was decreased (Figure 4C). However, no apparent differences in the levels of both precursor and mature forms of 23S rRNA were observed among WT, *atrsmD*, and *atrsmD pAtRsmD:AtRsmD-Myc*, as well as *atrimM*, strongly suggesting a specific role of m^2^G915 modification in the accumulation and maturation of the chloroplast 16S rRNA.

### AtRsmD and AtRimM are necessary for chloroplast translation and proteostasis

To further ensure that AtRimM is required to coordinate the AtRsmD-mediated 16S rRNA m^2^G915 modification *in planta*, a *var2-9 atrimM* double mutant was created, and its foliar phenotype was compared with *var2-9.* We found that loss of AtRimM substantially rescued *var2-9* phenotypes, including leaf variegation and total chlorophyll content (Figure 5A and B), similar to the impact of *AtRsmD* knockout in *var2* alleles (Figure 1G). Consistent with the plant phenotypes, the resulting immunoblot analyses found a significant decrease in the steady-state levels of chloroplast-encoded but not nuclear-encoded proteins in both *atrsmD* and *atrimM* mutants relative to WT (Figure 6A), indicating that 16S rRNA m^2^G915 modification is required to facilitate chloroplast translation. Note that the reduced chloroplast translation rescues the leaf variegation phenotype of *var2* by lowering the threshold level of the necessity of FtsH protease (69,70).

**Figure 5.**
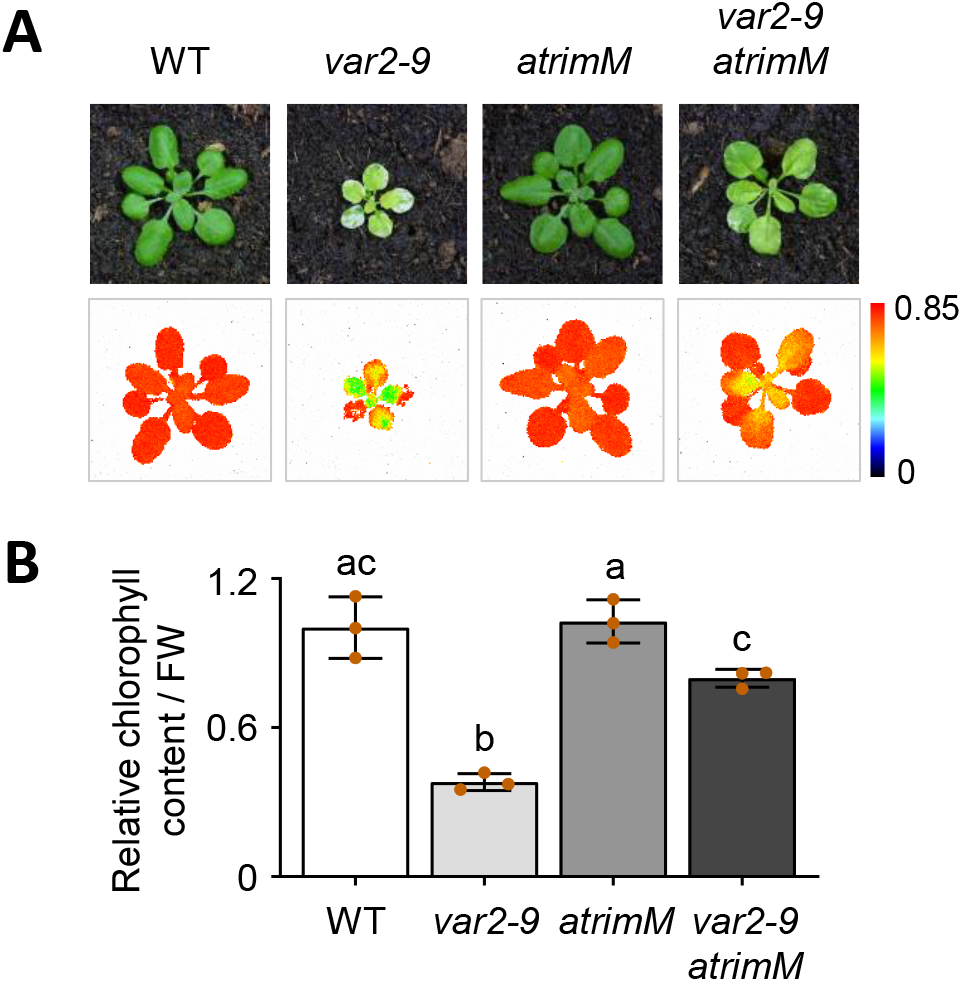
*AtRimM* mutation partially suppresses leaf variegation in *var2.* (**A**) and (**B**) Foliar phenotype (**A**, top panels) and Fv/Fm (**A**, bottom panels), and total chlorophyll content (**B**) were examined in 3-week-old WT, *var2-9, atrimM*, and *var2-9 atrimM* plants grown on soil under CL. More than 20 plants were used for each experiment, and the representative images are shown at the same scale (**A**). In (**B**), data are presented as mean ± SD (*n*=3), and lowercase letters indicate statistically significant differences between mean values (*P* < 0.01, one-way ANOVA with posthoc Tukey’s HSD test).

**Figure 6.**
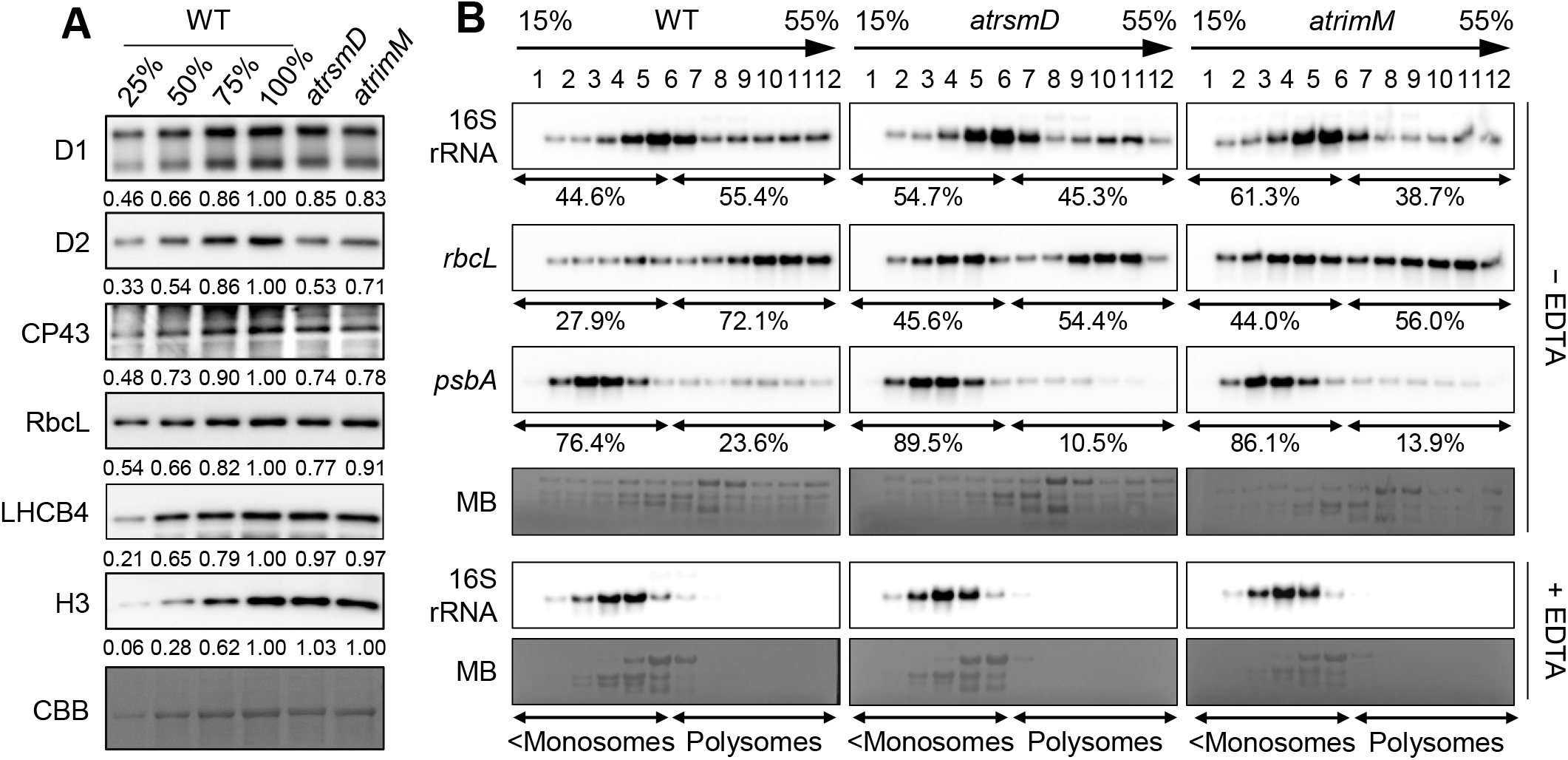
Loss of both AtRsmD and AtRimM reduces 16S rRNA levels in polysomes. (**A**) Immunoblot results showing the abundances of chloroplast-encoded proteins, such as PSII core proteins (D1, D2, and CP43) and nucleus-encoded chloroplast proteins, such as LHCB4 (a light-harvesting protein in PSII), in CL-grown 5-day-old seedlings of WT, *atrsmD*, and *atrimM.* Numbers at the bottom of each immunoblotting indicate the relative quantities of indicated proteins with or without serial dilution. Antibody against histone H3 and SDS-PAGE gel stained with CBB were used as loading controls. (**B**) Polysome loading of selected chloroplast RNAs in CL-grown 5-day-old seedlings of WT, *atrsmD*, and *atrimM.* Fractions from sucrose density gradients in the absence or presence of EDTA were analyzed by RNA gel blots using gene-specific probes. Lanes 1 to 12 indicate the fractions on the sucrose density gradients from the top (15%) to the bottom (55%). Signals in the polysomes (fractions 7-12) and monosomes/free RNAs (<Monosomes; fractions 1-6) fractions were quantified by the ImageJ software, and their relative quantities were indicated as percentage of the total signal over the 12 fractions.

To examine whether loss of either AtRsmD or AtRimM modulates the levels of 16S rRNA in polysomes, mRNA loading to polysomes, and ribosome assembly, we performed a sucrose density gradient fractionation followed by RNA gel blot analysis with WT, *atrsmD*, and *atrimM* seedlings. The RNA gel blot results showed that the levels of mature 16S rRNA in polysome fractions (fractions 7-12) were remarkably reduced in *atrsmD* and *atrimM* mutants compared to WT (Figure 6B), indicating both mutants contain fewer polysomes than WT. Moreover, chloroplast mRNAs, such as *rbcL* and *psbA*, were shifted towards the low sucrose density fractions in *atrsmD* and *atrimM* mutants compared to WT, which concurs with the reduced abundance of chloroplast proteins (Figure 6A).

## DISCUSSION

This study demonstrated the interrelated functionality of 16S rRNA m^2^G915 methyltransferase AtRsmD and ribosome assembly factor AtRimM for 16S rRNA maturation, ribosome assembly, and proteostasis in chloroplasts. The forward genetic screen of *var2* mutant lacking functional FtsH2 protease, involved in the degradation of damaged PSII core proteins, provided a clue that the plastid-specific AtRsmD plays a critical role in translation and proteostasis. Thus, its deficiency reduces the threshold level of conditional requirement of FtsH in the regulation of PSII proteostasis, thereby restoring leaf greening in *var2.* Therefore, this finding reveals the latent function of AtRsmD in chloroplasts under normal growth conditions, in addition to the earlier report that AtRsmD has a specific role in chloroplast translation under cold stress conditions (12).

Consistent with the impact of AtRsmD mutation in *var2*, significantly impaired m^2^G915 modification and maturation of 16S rRNA were found in the *atrsmD* mutant in the absence of external stimuli (Figure 4C; Supplementary Figure S4C and D). The impaired m^2^G915 modification specifically caused a significant maturation defect in 16S rRNA (Figure 4C), which decelerated protein translation, greening, and PSII activity (Figure 1G and 6). Chloroplasts also require the 16S rRNA m^4^C1352 methyltransferase CMAL for chloroplast ribosome biogenesis and plant development under normal growth conditions (23). Unlike AtRsmD, whose absence only impairs 16S rRNA maturation, the loss of CMAL disturbs the maturation of both 16S and 23S rRNAs, indicating its crucial function towards ribosome biogenesis. Consistent with the 16S and 23S rRNA maturation defect, *cmal* mutants show severe growth retardation. The impaired auxin synthesis and signaling were attributed as potential causes of the growth retardation in *cmal*, shedding new light on the link between ribosome biogenesis and auxin metabolism/signaling (23). Given that the *AtRsmD* mutation compromises chloroplast translation in young seedlings and greening in juvenile leaves (Figures 1G and 5A) and that the *atrsmD* mutant is fully viable despite the defect in 16S rRNA maturation (Figure 4C), it seems that AtRsmD functions only at the early stage of chloroplast biogenesis (Figure 1G) and under cold stress conditions (12). Therefore, we concluded that 16S rRNA m^2^G915 modification does not play a primary role but rather an auxiliary role, particularly during early chloroplast development that demands massive protein synthesis for a fast and accurate build-up of the entire photosynthetic apparatus. Likewise, AtRsmD may sustain 16S rRNA m^2^G915 modification in response to cold stress to reinforce chloroplast proteostasis.

Compelling evidence suggested that bacterial RsmD catalyzes the m^2^G966 modification on a fully assembled 30S ribosomal subunit (25). Our study on AtRsmD interactome and ribosome (monosome vs. polysome vs. polysome with EDTA) analyses also indicated an AtRsmD association with the 30S ribosomal subunit (Figure 2). Intriguingly, besides its interaction with ribosomal proteins, such as RPS13 (nuclear-encoded) and RPS19 (plastid-encoded), AtRsmD directly interacted with the nuclear-encoded ribosome maturation factor AtRimM in chloroplasts (Figure 2A to C; Supplementary Table S2). RimM functions in the assembly of the 30S ribosomal subunit, since RimM mutation in both bacteria and maize leads to a significant defect in 16S rRNA maturation (20,64). During the assembly of bacterial 30S subunit, RimM directly binds to RPS19 through its PRC β-barrel domain and facilitates the RPS19 association with the 16S rRNA, where helices 31 and 33b of 16S rRNA interact with RPS13 and RPS19, respectively, facilitating the interaction between RPS13 and RPS19 (64,71). It is important to note that the AtRsmD catalytic site G915 is located in the helix 31 of 16S rRNA and that RPS19 binding to the helix 33b of 16S rRNA is indispensable for RsmD binding to the 16S rRNA in bacteria (61). Therefore, it is rational to suggest that AtRimM may also serve as an additional key player in promoting 16S rRNA m^2^G915 modification by positioning AtRsmD near helix 31. Indeed, the primer extension-qPCR assay confirmed a substantial reduction of 16S rRNA m^2^G915 modification in the *atrimM* mutant relative to WT seedlings. The suppression of leaf variegation in *var2* mutant by the loss of AtRimM also supports the conclusion that AtRimM is required for the AtRsmD-dependent 16S rRNA m^2^G915 modification and chloroplast proteostasis.

In summary, this study reveals that AtRsmD methyltransferase and ribosome assembly factor AtRimM both function in 16S rRNA maturation and ribosome assembly, which is critical for sustaining chloroplast proteostasis during early chloroplast biogenesis. We also believe this finding will stimulate studies on delineating the process of chloroplast ribosome assembly and finding critical factors involved in the spatiotemporal regulation of 16S rRNA methylation in plants.

## ACKNOWLEDGEMENTS

We thank the Core Facility of Genomics and Proteomics in the Shanghai Center for Plant Stress Biology (PSC) for performing RNA sequencing and mass spectrometry, respectively.

