## Supplemental Figures 1-5 for "Plastid-specific RsmD methyltransferase and ribosome maturation factor RimM are crucial for 16S rRNA maturation and proteostasis"

### Supplementary Figure S1

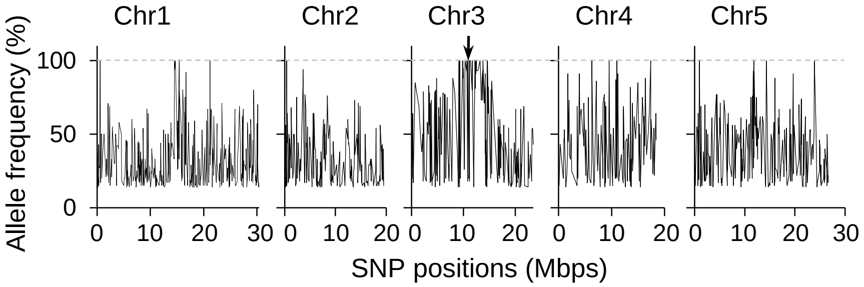


**Supplementary Figure S1.** Whole genome sequencing was performed (See MATERIALS AND METHODS) to identify a causative mutation of *svr12*. The result shows the SHOREmap analysis of SNP concurrence across the five Arabidopsis chromosomes from a pooled mapping population of *svr12*. An enrichment of SNP concurrence can be seen in chromosome 3, where *AtRsmD* (At3g28460; encoding chloroplast RsmD methyltransferase) is located, as indicated by the arrow.

### Supplementary Figure S2

```
AtRsmD        114 LQVLAGTAKRKKLLSLKGMDVRPMMEVVKGAAFGILQAAGGCPTSLRPGRWLDLYSGTGS 173
P10120         12 IRIIGGQWRGRKLPVPDSPGLRPTTDRVRETLFNWLAP------VIVDAQCLDCFAGSGA 65
WP_005693161   13 VRIIAGLWRGRKLPVLNSEGLRPTGDRVKETLFNWLMP------YIHQSECLDGFAGSGS 66
CAB11354        1 MRVISGSKKGRSLKAVAGTSTRPTTDKVKESIFNMIGP------YFDGGRGLDLFAGSGG 54
BAM54557        5 MRIY----GNRLLKTLPGQQTRPTTGKVRLALFNIWRG------EIQHCHWLDLCAGNGT 54
AAA50947       28 LDPDHRRCRRRRRIAVPPRGTRPTTDRVRESLFNIVTARR----DLTGLAVLDLYAGSGA 83
WP_002656751   13 -YVSSGKYKGKKILFPKNGSVRPVMSLVREAFFSIIFK------DIVNSKFLDVFAGTGI 65
WP_001132760   10 FKIIGGACKGLGLNLPNISSTRPTKAIVRESFFNTLQA------EINGAHFIEVFSGSAS 63
WP_062431539    1 MRIY----GNRQIKTLPGELTRPTLAKVREAIFNIWQG------QVSGCRWLDLCAGSGS 50
WP_009554370    3 LRIY----GNRALKTLPGRDTRPTLARVREAVFNIWQG------KIEGCRWLDLCAGTGS 52

AtRsmD        174 VGIEAISRGCSEAHFVEMDPWVVSNVLQPNLEHTGFVDASVIHTARVENFLER----ADK 229
P10120         66 LGLEALSRYAAGATLIEMDRAVSQ-QLIKNLATLKAG-------N--ARVVNSNAMSFLA 115
WP_005693161   67 LGFEALSRQAKKVTFLELDKTVAN-QLKKNLQTLKCS-------SEQAEVINQSSLDFLK 118
CAB11354       55 LGIEALSRGFEHCIFVDRDFKAIQ-TVKSNLKTLELT-------KHAQVYRNDAERALHA 106
BAM54557       55 LGAEALCRDADKVVAIEQSAKVCT-IIKENWQKLAKP-------GQQWSVLRGDVLKLL- 105
AAA50947       84 LGLEALSRGAASVLFVESDQRSAA-VIARNIEALGLS-------G--ATLRRGAVAAVV- 132
WP_002656751   66 MSVEALSRGASLAHLVECNRKIKI-TLVENFSFVEEF----------YKFFFQRAEDFL- 113
WP_001132760   64 MGLEALSRGAKSAVFFEQNKSAYK-TLLENISLFKNR---LKKEMEIQTFLDDAFKLLPT 119
WP_062431539   51 MGAEALCRGAVKVVGIEKNSQACR-IIQENWQKVAKV-------DQEYQILKGDLLKRV- 101
WP_009554370   53 MGAEALCRGAAIAIAVEKSSRACA-VIQQNWQQVART-------DQTFQVLRGDVVKRL- 103

AtRsmD        230 LVGKDGVFDYISVTPPYMEVDYE----VLMDQIAK---SPAIGENTFILVEYPSRTTMLD 282
P10120        116 QK--GTPHNIVFVDPPFRRGLL----EETINLLED--NG-WLADEALIYVESEVENGLPT 166
WP_005693161  119 QPQNQPHFDVVFLDPPFHFNLA----EQAISLLCE--NN-WLKPNALIYVETEKDKPL-I 170
CAB11354      107 AAKRETGFRGIFLDPPYKEQKL----KALLTLIDE--YQ-MLEEDGFIVAEHDREVELPE 159
BAM54557      106 PSLAGQTFDRIYFDPPYGSGLY----NPVLTLVGE--LQ-ILSPTGEMAVEYDRHHWQPP 158
AAA50947      133 AAGTTSPVDLVLADPPYNVDSADV--DAILAALGT--NG-WTREGTVAVVER-ATTCAPL 186
WP_002656751  114 -SKKDLFYDFIYLDPPFNYKNKI-------NLLEIILKGKILNDKVSIIMHYPSNEDLEI 165
WP_001132760  120 LCLKNGVLNIIYLDPPFETSGFLGIYEKCFQALERLLKR-FNPKNLLVVFEHESMHEMPK 178
WP_062431539  102 ENLGGETFDLIYFDPPYAAKLY----DRVLAKIVD--LE-LLAPSGELAVEYDPKLWQPL 154
WP_009554370  104 PTLAGQTFDRIYFDPPYAEDLY----QPVLDEIAS--YA-LLASDGELAVEHSPDRTDII 156

AtRsmD        283 S-CGCLEKMTDRRFGRTHLAIY 303 Arabidopsis thaliana
P10120        167 V-PANWSLHREKVAGQVAYRLY 187 Escherichia coli str. K-12 substr. MG1655
WP_005693161  171 T-PENWTLLKEKTTGIVSYRLY 191 Haemophilus influenzae
CAB11354      160 T-VEIL---------------- 164 Bacillus subtilis subsp. subtilis str. 168
BAM54557      159 EQIAGLELMRQKKYGLSNLAFY 180 Bacillus subtilis BEST7613
AAA50947      187 TWPEGWRRWPQRVYGDTRLELA 208 Mycobacterium tuberculosis
WP_002656751  166 N-TSKFSVYNLKRYGGSKLIFL 186 Borreliella burgdorferi
WP_001132760  179 S-LVTLAIIKQKKFGKTTLTYF 199 Helicobacter pylori
WP_062431539  155 E-LPGLELFKEKNYGKTAIAFY 175 Synechococcus sp. PCC 73109
WP_009554370  157 T-PPSLEICRQKAYGNSALTFF 177 Oscillatoriales cyanobacterium JSC-12
```

**Supplementary Figure S2.** Multiple sequence alignment of AtRsmD and representative bacterial RsmD proteins. Black and grey boxes indicate identical and similar amino acids, respectively. The conserved amino acid sequences were aligned by using the ClustalW program before shading with the Boxshade program.

### Supplementary Figure S3

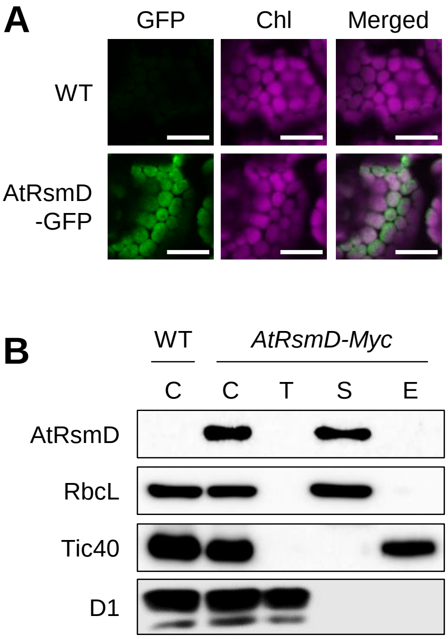


**Supplementary Figure S3.** AtRsmD resides in the chloroplast stroma. (**A**) Subcellular localization of AtRsmD was examined by confocal microscopy using 5-day-old transgenic *svr12* seedlings expressing *AtRsmD-GFP* under the control of CaMV 35S promoter (*p35S:AtRsmD-GFP*). All images were taken at the same scale. Scale bars: 10 µm. Chl, chlorophyll. (**B**) Chloroplast subfractionation and immunoblot analyses. Total chloroplasts (C) isolated from 3-week-old *atrsmD pAtRsmD:AttRsmD-Myc* plants were subjected to sucrose density gradient centrifugation to isolate thylakoid (T), stroma (S), or envelope (E) fractions. Antibodies against proteins localized in thylakoid (D1), stroma (RbcL), and inner envelope membrane (Tic40) were used to verify the purity of the fractions.

### Supplementary Figure S4

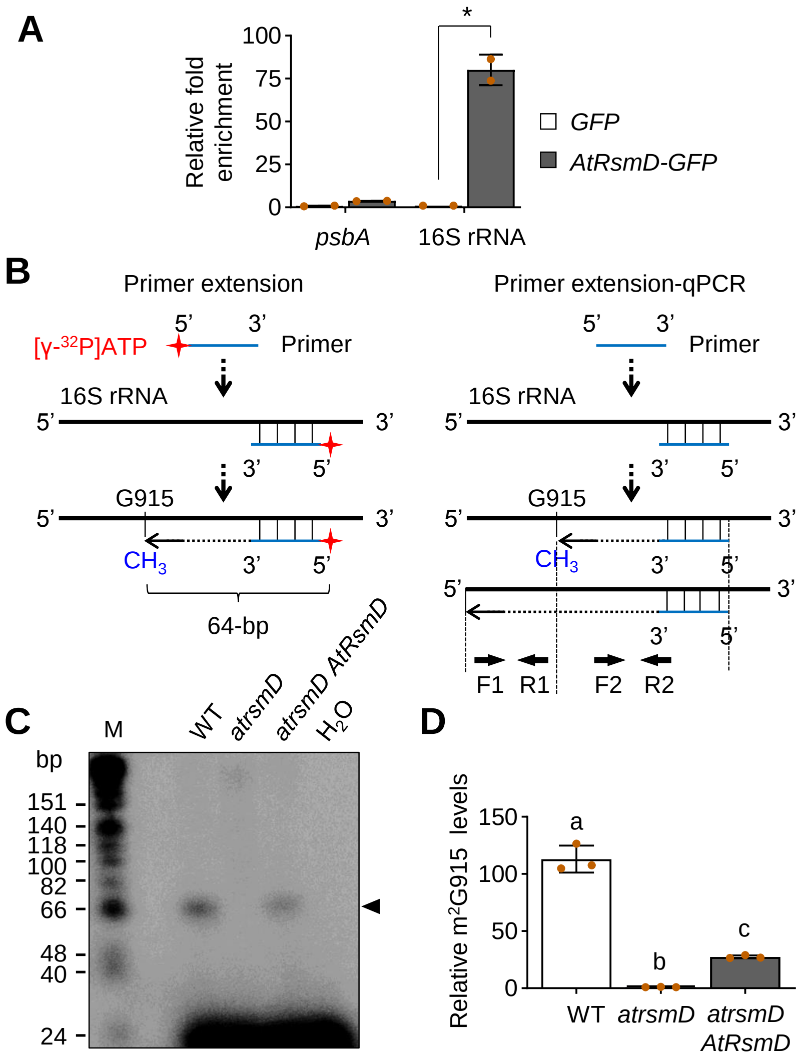

**Supplementary Figure S4.** AtRsmD directly binds chloroplast 16S rRNA to catalyze $m^2G915$ modification. (**A**) RNA-immunoprecipitation (RIP) coupled with RT-qPCR analyses using 5-day-old seedlings of *svr12 p35S:AtRsmD-GFP* (*AtRsmD-GFP*) and WT expressing free GFP (GFP). RNA bound to AtRsmD-GFP were immunoprecipitated with a GFP-trap bead, and the input and eluate were used for RT-qPCR analysis to examine AtRsmD binding to chloroplast 16S rRNA. Chloroplast-transcribed *psbA* was used as a negative control. The enrichment value was normalized to the input, representing means ± SD ($n$=2). Asterisk indicates statistically significant differences between mean values by Student's *t*-test (P < 0.01). (**B**) Schematic diagrams for experimental procedures of primer extension (left side) and primer extension coupled to qPCR (primer extension-qPCR; right side) assays to detect or quantify the $m^2G915$ modification. For the primer extension assay, the chloroplast 16S rRNAs were reverse-transcribed using a gene-specific primer end-labeled with [γ-$^{32}$P]ATP. The primer was designed to generate a 64-base pair (bp) product with $m^2G915$ modification. To quantify the level of $m^2G915$ modification, primer extension products reverse-transcribed with a primer, i.e., reverse-complementary to the 16S rRNA nucleotides (1092-1108; downstream of the G915), were used for the qPCR. As indicated, the qPCR primer pair F1 and R1 was designed to amplify the reverse-transcribed 16S rRNA with G915, while the F2 and R2 primer pair was used to amplify the 16S rRNA pool regardless of $m^2G915$ modification. The relative level of $m^2G915$ modification was determined by quantifying non-methylated 16S rRNA G915 (F1/R1) versus total amount of 16S rRNA (F2/R2). (**C**) and (**D**) Primer extension (**C**) and primer extension-qPCR (**D**) assays using total RNA extracted from CL-grown 5-day-old seedlings of WT, *atrsmD*, and *atrsmD pAtRsmD:AtRsmD-Myc* (*atrsmD AtRsmD*) were performed as described in (**B**). The $^{32}$P-labeled primer-driven extension products were subjected to a denaturing polyacrylamide gel and visualized by phosphorimaging (**C**). In parallel, the reaction was conducted without RNA ($H_2O$). The arrowhead indicates the extension products containing $m^2G915$. M: size marker. In (**D**), data are presented as mean ± SD ($n$=3), and lowercase letters indicate statistically significant differences between mean values ($P < 0.01$, one-way ANOVA with posthoc Tukey's HSD test).

### Supplementary Figure S5

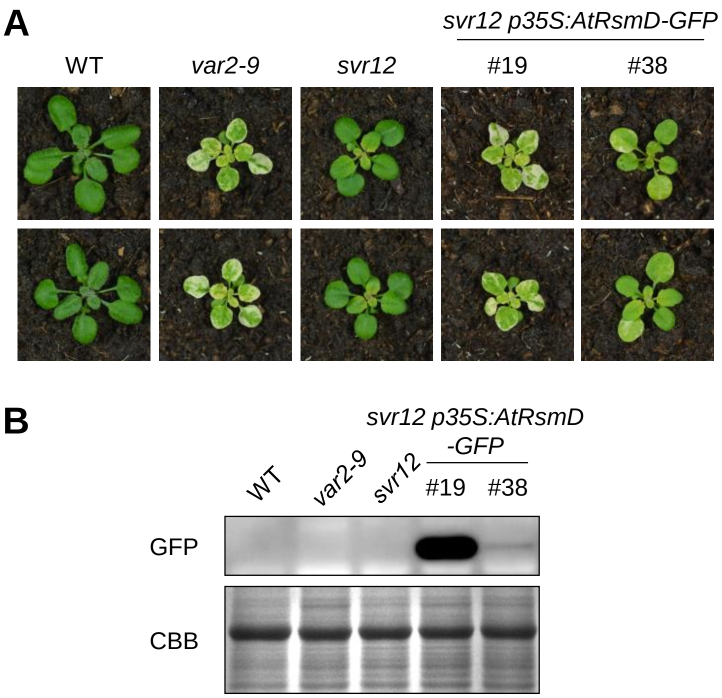


**Supplementary Figure S5.** Complementation assay by expressing AtRsmD-GFP in *svr12*. (**A**) and (**B**) Foliar phenotype of two different plants per line (**A**) and protein abundance of AtRsmD-GFP (**B**) were examined in CL-grown 3-week-old WT, *var2-9*, *svr12*, and two independent *svr12 p35S:AtRsmD-GFP* transgenic plants. The representative images are shown at the same scale (**A**). GFP antibody was used for the immunoblot analysis, and SDS-PAGE gel stained with CBB was used as a loading control (**B**).
